# Denaturing purifications support direct interaction between PRC2 and RNA in cells

**DOI:** 10.64898/2026.05.19.725914

**Authors:** Stephen Henderson, Lucia Conde, Alex Hall Hickman, Samuel Marguerat, Richard G. Jenner

## Abstract

Polycomb Repressive Complex 2 (PRC2) maintains repression of genes specific for other cell differentiation states. PRC2 binds RNA in vitro with a preference for G-rich sequences. UV-based crosslinking coupled with immunoprecipitation (CLIP) experiments have shown that PRC2 also binds RNA in cells. Recently, Guo et al reported that a stringent denaturing variant of CLIP called CLAP did not detect PRC2 RNA binding in cells. We present a reanalysis of CLAP data that supports direct interaction of PRC2 with RNA in cells. CLAP for Halo-tagged PRC2 subunits from mixed populations of human and mouse cells specifically enriched for RNA from the species in which the proteins were tagged. The lack of apparent PRC2 RNA binding in Guo and colleagues’ analysis stems from a scaling step that deflates enrichment scores for low-complexity CLAP samples. Our findings pave the way for studies seeking to determine the physiological roles of PRC2 RNA binding activity.

---

PRC2 modifies chromatin to maintain genes specific for other cell types and differentiation stages in a repressed state. PRC2 is essential for cell differentiation, from the earliest stages of embryogenesis through to adulthood.

PRC2 comprises a catalytic core made up of EZH1 or EZH2, EED, and SUZ12. EZH1 and EZH2 methylate histone H3 at lysine 27 (H3K27me3), which in turn recruits the canonical forms of PRC1 via their CBX subunits (reviewed in ^1^). The association of PRC2 with various accessory factors defines two sub-forms of the complex that differ in their mode of recruitment to chromatin: PRC2.1 is recruited to CpG islands, while PRC2.2 is recruited to H2AK119ub deposited by variant forms of PRC1.

PRC2 has long served as a paradigm for chromatin regulatory proteins that also bind RNA. Mediated by its catalytic core, PRC2 preferentially interacts with G-quadruplex RNA structures, exhibiting an affinity on the order of 10-100 nM ^2–7^. The use of UV irradiation to covalently crosslink proteins to the RNAs with which they directly interact demonstrates that PRC2 binds nascent RNAs in cells and confirms the preference of the complex for G-rich sequences ^4,6,8,9^.

In vitro, RNA inhibits PRC2 catalytic activity and interaction with DNA and nucleosomes ^2,4,5,9–12^. Cryo-EM has revealed that G4 RNA induces PRC2 dimerization, internalising the active site in an inaccessible position, and thus providing a mechanism for the inhibitory effect of G4 RNA ^13^.

RNA also inhibits PRC2 function in cells. Depleting RNA from cells increases PRC2 chromatin association ^9,14^ while ectopically tethering G4-forming RNAs to repressed genes removes PRC2 from chromatin and reduces H3K27me3 in cis ^4,7^. Residues in PRC2 that mediate its RNA binding activity have been identified but the importance of these residues for nucleosome binding has precluded the generation of separation of function mutants necessary to test the biological importance of PRC2 RNA binding in cells ^15,16^.

Seemingly conflicting with these data, Guo and colleagues recently reported that PRC2 does not bind RNA in living cells ^17,18^. CLIP of v5-tagged EZH2, EED or SUZ12 from a mixture of crosslinked human cells expressing the tagged subunit and crosslinked mouse cells without the tag identified both human and mouse RNAs bound by PRC2, demonstrating that the method can produce false positive results. Furthermore, interrogation of PRC2 RNA binding in cells using CLAP (covalent linkage and affinity purification), a variant of CLIP that can be performed under fully denaturing conditions due to covalent interactions formed between HaloTag proteins and HaloLink resin, was reported not to identify any PRC2-bound RNAs but was able to detect RNAs bound by the canonical RNA binding proteins PTBP1 and SAF-A (HNRNPU). Based on these data, the authors concluded that although PRC2 has the capacity to bind RNA in vitro, it does not do so in cells, potentially due to compartmentalisation or competition with other proteins that bind RNA with higher affinity. We and others have noted that UV-crosslinking methods favour detection of protein-RNA interactions that involve amino acids often found in canonical RNA binding domains, such as RNA recognition motifs, but not necessarily in other types of RNA binding surfaces, such as those present on PRC2 ^7,19^.

We sought to confirm whether or not CLAP methodology supported the conclusion that PRC2 does not bind RNA in cells. We processed the CLAP data from Guo and colleagues following the analysis framework described by the authors and using an alternative approach and report that both schemes support direct interaction between PRC2 and RNA in cells.

## RESULTS

### The analysis framework of Guo et al demonstrates PRC2 RNA binding in cells

Guo and colleagues performed CLAP for the core PRC2 subunits EED, EZH2 and SUZ12 ^17^. To allow distinction between genuine and spurious RNA binding events, crosslinked human HEK293 cells expressing a HaloTag version of the subunit of interest were mixed with crosslinked unmodified mouse embryonic stem cells (“+tag” samples), with the expectation that if PRC2 binds RNAs in cells then only human RNAs should be identified as enriched relative to input RNA. Any enriched mouse RNAs must instead represent interactions with PRC2 that have occurred after cell mixing and lysis. Reciprocally, “-tag” samples of mouse cells expressing a HaloTag form of the protein of interest mixed with unmodified human cells were also prepared. In addition, a CLAP experiment was performed for GFP, which is not expected to bind RNA in cells, and for the canonical RNA binding proteins SAF-A and PTBP1, which served as positive controls.

We first sought to analyse the data according to the framework described by Guo and colleagues. Reads aligning to a set of multi-copy RNAs were removed and the remaining reads mapped to the combined human and mouse genomes. Remaining reads mapping to multi-copy RNA loci in the genome were then removed in a second filtering step. Genes were divided into 100 bp windows, and the number of reads in each window counted for each sample. Bound RNA regions were then identified as windows in which there were significantly more reads in the CLAP sample versus the respective input sample.

A key aspect of Guo and colleagues’ reported analysis framework is that the number of reads mapping to each gene window is normalised by the number of reads mapping to all RNAs ^17,18^. In this way, for an RNA to be identified as bound by the protein of interest it must be enriched relative to all other RNAs in the cell, including multi-copy RNAs, and relative to the input sample.

Following these analysis steps, we observed significant enrichment (p< 10^-6^) of large numbers (tens of thousands) of human RNA regions in SAF-A and PTBP1 +tag CLAP experiments, confirming that these proteins directly bind RNA in cells (Fig 1A). In contrast, only a handful of human RNA regions were enriched in the GFP +tag sample or the respective −tag control samples (Fig 1B), demonstrating minimal background RNA pull-down.

**Fig. 1.**
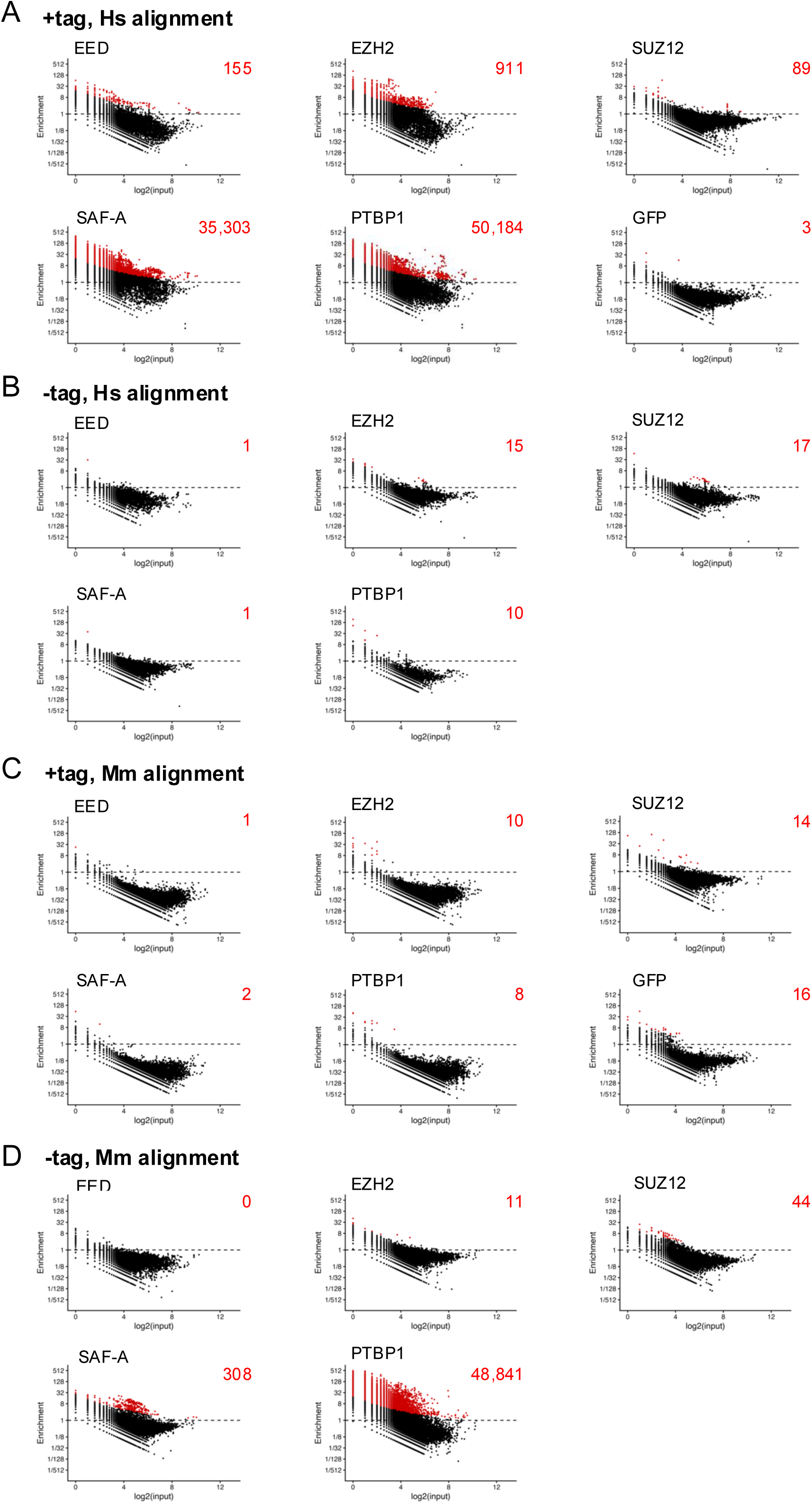
The analysis framework of Guo et al demonstrates PRC2 RNA binding in cells. A. Scatterplots of CLAP enrichment ratio versus paired input read count (both log values) for +tag samples within 100nt windows across human genes. Windows with significant enrichment (p <10^−6^; binomial test) are shown in red and enumerated in the top right of each plot. B. As A., except for −tag CLAP control samples. C. As A., except for +tag CLAP samples and mouse genes. D. As A., except for −tag CLAP samples and mouse genes.

Next, turning our attention to PRC2, we observed enrichment of 911 human RNA windows in the EZH2 +tag CLAP experiment (Fig 1A). Although lower than for SAF-A and PTBP1, the number of enriched RNA windows was orders of magnitude higher than for the control GFP +tag dataset (3 windows) and −tag control sample (15 windows; Fig 1B). We also observed some enrichment of human RNA windows in EED and SUZ12 +tag experiments, albeit fewer than for EZH2, consistent with structural data ^13^. Over 50% of windows significantly enriched by EED CLAP were common with EZH2, with SUZ12 instead showing very few enriched windows in common (Fig S1). The specific enrichment of RNA windows in PRC2 +tag experiments and not in controls was also apparent at the lower p<10^-5^ threshold and thus not dependent on a particular p-value cut-off (Fig S2).

The enrichment of human RNAs in +tag but not −tag PRC2 CLAP samples indicated detection of PRC2 crosslinking to these RNAs in cells prior to cell mixing and lysis. If this was the case, we reasoned that crosslinking to mouse RNAs should show a reciprocal pattern, with stronger signals in −tag samples compared to their respective +tag samples. This pattern was indeed evident for SAF-A and PTBP1, with each −tag sample showing significant enrichment of mouse RNA windows relative to input that was more extensive than for their respective +tag control experiments (Figs 1C and D). Thus, these data support both of these proteins directly binding to RNA in living mouse cells. However, the detection of SAF-A RNA binding in mouse cells was over 100-fold reduced compared to human cells (compare −tag Mm alignment (Fig 1D) with +tag Hs alignment (Fig 1A)), suggesting reduced crosslinking efficiency in the mouse cells.

EED, EZH2 and SUZ12 −tag samples did not exhibit significant enrichment of mouse RNA regions relative to input. However, given the much weaker detection of SAF-A RNA binding in mouse versus human cells, the lack of significant PRC2 RNA binding events in mouse could be due to low sensitivity. Indeed, although not significant at p<10^-6^, the enrichment scores for mouse RNA windows were greater (shifted upwards) in the EED and EZH2 −tag samples (Fig 1D) compared to their respective +tag samples (Fig 1C). SUZ12, by contrast, exhibited similar enrichment scores for mouse RNA windows in +tag and −tag samples.

To guard against the results reflecting anomalous PRC2 input samples, and because all input samples should mostly exhibit the same level of RNAs, we also normalised the +tag CLAP data against a single merged +tag input and normalised −tag CLAP samples against a single merged −tag input. This analysis maintained the stronger enrichment of human RNA in PRC2 +tag versus control experiments, while increasing the number of enriched windows, likely due to the increased statistical power (Fig S3).

Thus, from our analysis of Guo and colleagues’ CLAP data we conclude: 1) human RNAs are significantly enriched in EZH2 and, to a lesser extent, EED and SUZ12 +tag CLAP experiments compared to input; 2) human RNAs are enriched in PRC2 +tag CLAP experiments compared to GFP and −tag control samples; and 3) for EZH2 and EED, mouse RNAs are correspondingly enriched in −tag samples versus +tag control samples. These findings are consistent with direct binding of PRC2 to RNA in living cells.

### Data scaling by the proportion of genome-mapped reads accounts for the lack of detection of PRC2 RNA binding by Guo et al

We considered why we observed significant enrichment of human RNA windows in PRC2 CLAP samples versus input while Guo and colleagues did not. The RNA enrichment scores we calculated for EED, EZH2 and SUZ12 were higher than those calculated by Guo et al, while the scores we calculated for SAF-A and PTBP1 were lower (Fig 2A). To identify the source of these differences, we examined the data at each stage of the analysis process.

**Fig. 2.**
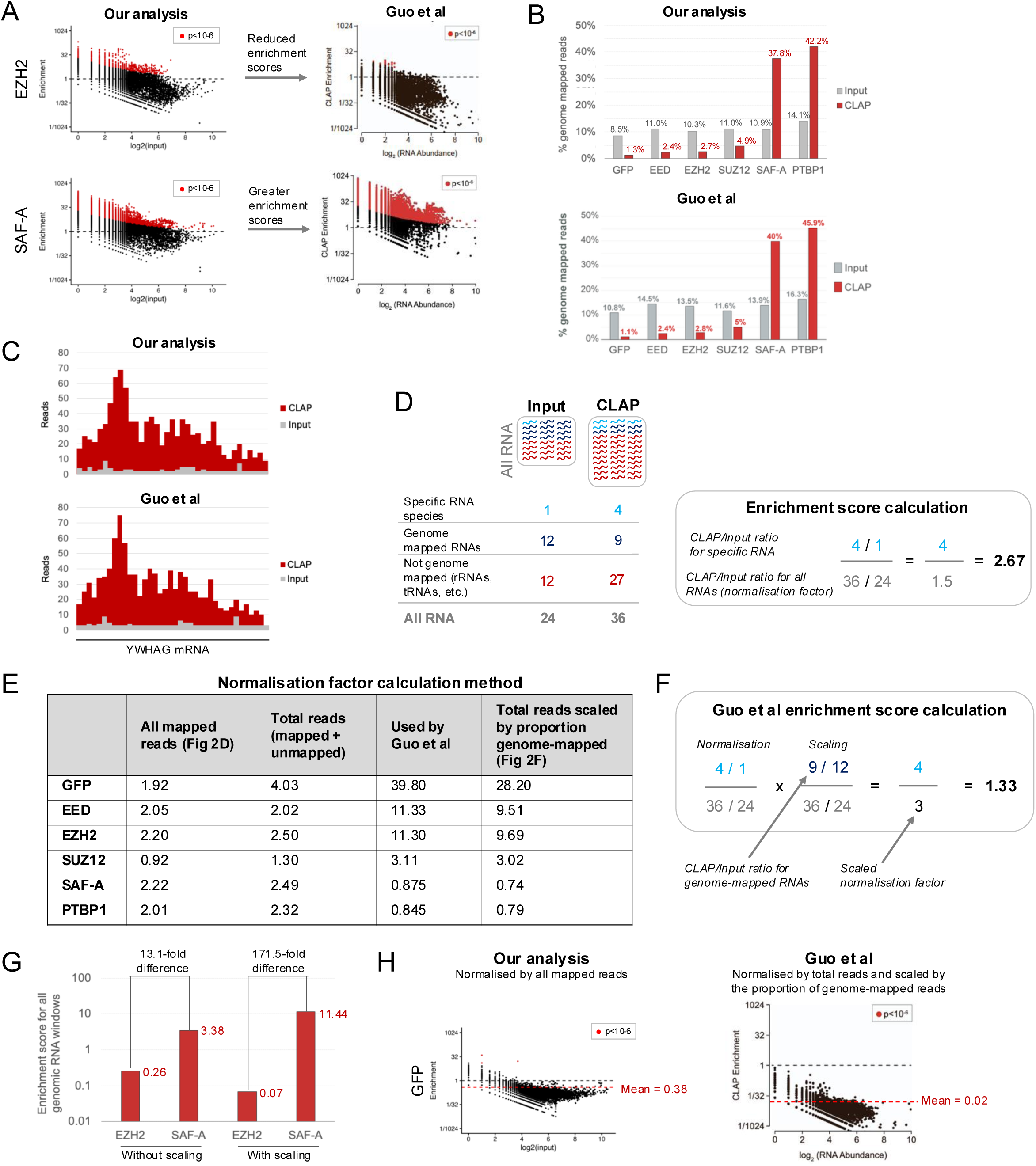
An additional scaling step accounts for lack of detection of PRC2 RNA binding by Guo et al. A. Comparison of enrichment scores from our and Guo and colleagues’ analyses. EZH2 and other PRC2 subunits exhibit reduced enrichment scores (y-axis) in Guo’s analysis, while SAF-A and PTBP1 exhibit increased enrichment scores. B. Proportions of reads from +tag CLAP and matched input datasets mapped to the human + mouse genomes (after removal of reads mapping to multi-copy RNAs) in our analysis (top) and Guo’s analysis (below; figure from ^18^). C. Read counts per window across human YWHAG mRNA in EZH2 +tag CLAP and input samples in our analysis (top) and Guo’s analysis ( below; data from GSE253466). D. Data normalisation methodology followed by this study and presented by Guo et al ^18^. The enrichment score for an RNA window is calculated as the ratio of CLAP vs input reads for that window, divided (normalised) by the ratio of all mapped CLAP vs input reads. E. Normalisation factors calculated using different methods. From left to right: using all mapped reads (as shown in D and used in this study), using total reads (mapped + unmapped), the factors used by Guo et al in their study, normalisation factors calculated using the scaling method shown in F. F. Data normalisation and scaling methodology that Guo et al appear to have followed. In addition to normalizing the data by the total number of reads, the data were also scaled by the proportion of reads mapping to the genome. G. Effect of data scaling on the enrichment scores for all genomic RNA windows for EZH2 and SAF -A. The difference between the enrichment scores is exaggerated by the scaling step. H. Effect of the additional scaling step on the apparent enrichment of RNA in the negative control GFP +tag CLAP dataset, which theoretically should exhibit an RNA profile similar to input RNA (mean enrichment score close to 1). The additional scaling step shifts th e mean enrichment score well below 1.

The proportions of reads mapping to the genome were similar between our and Guo and colleagues’ analyses, demonstrating that the difference did not stem from the alignment stage (Fig 2B). The numbers of reads mapping to windows within genes was also similar, so the difference did not lie at the read assignment stage (Fig 2C; see Methods for reasons why the numbers are not identical).

This suggested a difference in the downstream step of data normalisation. We had normalised read counts within each window by the total number of mapped reads (reads mapping to multi-copy RNAs or to the human or mouse genomes). This matches the procedure laid out by Guo and colleagues ^18^, which emphasised the importance of ensuring that the enrichment of specific RNA segments is relative to all the RNAs in an experiment (Fig 2D). This contrasts with the process followed by a study from Lee and colleagues that normalised the CLAP data by the number of genome-mapped reads ^23^.

The normalisation factors used by Guo et al can be derived from their supplemental data tables, which contain CLAP and input read counts along with normalised enrichment scores for each gene window. Comparison of the normalisation factors we calculated (Fig 2E, column 2) with those used by Guo and colleagues (Fig 2E, column 4) revealed clear differences, with the factors used by Guo having the effect of down-weighting the enrichment scores for PRC2 subunits and up-weighting the enrichment scores for SAF-A and PTBP1. Including or excluding unmapped reads made little difference to the normalisation (Fig 2E, column 3) and so we could exclude potential variations in read mapping from the differences in normalisation factors between our studies.

We instead considered that the difference may reflect a second normalisation step performed by Guo et al that was referred to in the methods section but not included in the normalisation method visualisations or accompanying rationalisation text ^18^. Specifically, the authors referred to accounting for differences in overall complexity between samples by scaling the total number of sequenced reads by the proportion of genome-mapped reads within each sample (Fig 2F). Adding this scaling step to the analysis produced normalisation factors similar to those used by Guo et al, which have the effect of reducing PRC2 enrichment ratios and increasing SAF-A and PTBP1 enrichment ratios (Fig 2E, last column, and Fig S4).

The justification for this scaling step is unclear. Low complexity datasets (those with a low proportion of genome-mapped reads) arise from amplification of low input samples. Such datasets have, by definition, correspondingly lower numbers of reads mapping to single-copy RNA windows, and the scaling step employed by Guo et al exacerbates, rather than compensates for, this difference. The effect of the scaling process can be visualised by considering its effect on all genomic RNA windows together in a low complexity dataset (EZH2) and a high complexity dataset (SAF-A) (Fig 2G). After normalisation without the scaling step, EZH2 exhibited an enrichment score for all genomic RNA windows of 0.26, while SAF-A exhibited an enrichment score for all genomic RNA windows of 3.38, a 13-fold difference. After the additional scaling step, the enrichment score for all genomic RNA windows for EZH2 dropped to 0.07 (0.26 * 0.26), but for SAF-A it increased to 11.44 (3.38 * 3.38), a 172-fold difference.

This effect of the scaling step is also apparent when considering the negative control GFP +tag dataset. Given that pull-down of GFP should not enrich for any particular RNA species, the read distribution should approximate those of input samples and thus the mean enrichment score should be close to 1. Normalising by the total number of mapped reads alone (our method) produced a mean log2 enrichment score of -1.41 (enrichment score of 0.38), while the additional scaling step performed by Guo and colleagues resulted in a substantially lower mean enrichment score of -5.51 (enrichment score of 0.02). Thus, compared to the scaling method of Guo et al, our normalisation method resulted in a read count distribution more closely matching that expected of a control CLAP experiment.

We conclude that the seeming lack of evidence for PRC2 RNA binding in the CLAP data analysis of Guo et al was due to the application of an additional scaling step that down-weights read counts from datasets with low proportions of genome-mapped reads.

### CLAP data analysis with an alternative method supports PRC2 RNA binding in cells

Given the potential issues surrounding Guo and colleagues’ data scaling methodology, we sought another means to quantify the enrichment of RNAs in CLAP samples. The presence of cells from a second species in which the protein of interest was not tagged provided a set of control RNAs against which the enrichment of experimental RNAs can be judged.

To do this, we used standard tools to calculate CLAP and input read counts per million mapped reads (all reads mapping to multi-copy RNAs or the human or mouse genomes) and further normalised CLAP sample read counts to their respective inputs. We then calculated the average input normalised read count for each human exon and intron in +tag and −tag samples. We found that human exons and introns tended to have higher read counts in the EED, EZH2 and SUZ12 +tag samples than in the control GFP +tag sample (Fig 3A). Similarly, human exons and introns displayed higher read counts in +tag samples than in their corresponding −tag controls (Fig 3B). These effects were most clearly seen for EZH2 but was also apparent for EED. Reciprocally, mouse exons and introns displayed higher read counts in EED and EZH2 −tag samples, in which the proteins are epitope tagged in mouse cells, than in +tag samples (Fig 3C). SUZ12 did not show this effect, consistent with the results of analysis with Guo and colleague’s statistical framework (Fig 1). The read counts for the canonical RNA binding proteins SAF-A and PTBP1 followed the same pattern as those of EZH2 and EED. Thus, these data confirm that CLAP for EED and EZH2 specifically enriched for RNA from the cells in which the protein was tagged, demonstrating direct binding to RNA in cells.

**Fig. 3.**
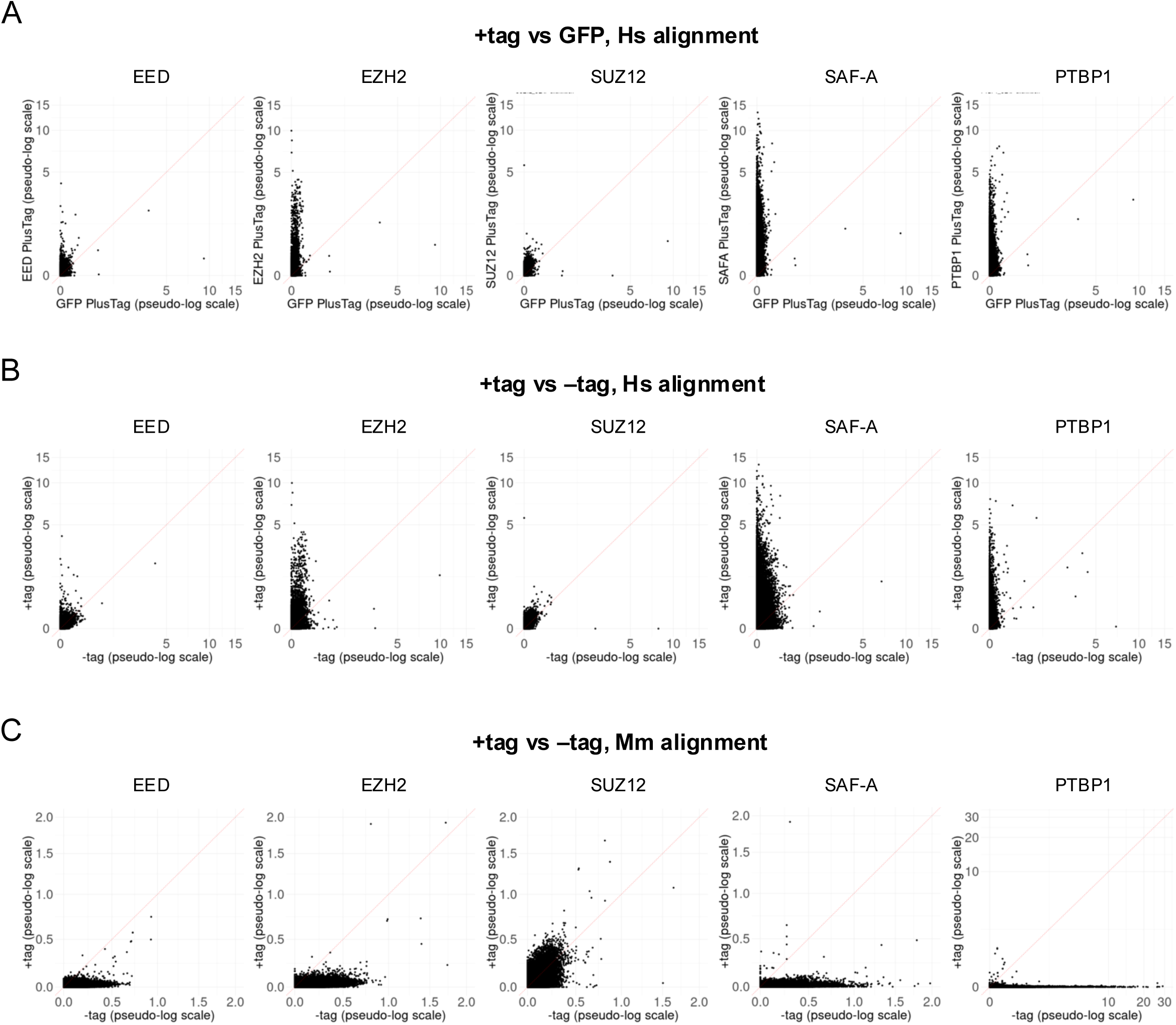
CLAP data analysis with an alternative method supports PRC2 RNA binding in cells. A. EED, EZH2, SUZ12, SAF-A and PTBP1 crosslinking to human RNA (input-normalised counts per million) in +tag samples (y-axis) versus the GFP +tag sample (x-axis). Each point represents an individual exon or intron. Data are shown on pseudo -log scales. The dotted red lines denote y=x. B. As A, except showing EED, EZH2, SUZ12, SAF-A and PTBP1 crosslinking to human RNA (input-normalised counts per million) in +tag samples (y-axis) versus −tag samples (x-axis). C. As B. except for mouse RNA. Note that the PTBP1 plot axis maxima differ from the other plots in the set.

### Visualising crosslinked RNA at individual genes supports PRC2 RNA binding in cells

We next asked if PRC2 RNA binding could be observed at individual genes. We first examined the human long non-coding RNA XIST that was the focus of the study by Guo and colleagues^17^. Our analysis reproduced the SAF-A and PTBP1 binding profiles but also demonstrated significant peaks (p< 10^-6^) of EED and SUZ12 binding at the 5’-end of the RNA that were not present in GFP or −tag controls (Fig S5). However, the peak was not well defined in the EZH2 +tag dataset.

We also visualised CLAP data after processing with standard tools to give raw counts per million mapped reads (genome-mapped + multi-copy RNA-mapped; Fig 4). Similar to the enrichment calculation using Guo’s method, we found that EED and SUZ12, but not EZH2, exhibited high signals at the 5’-end of XIST RNA that were not present in the GFP or −tag controls. These peaks also remained after normalisation to their respective input samples.

**Fig. 4.**
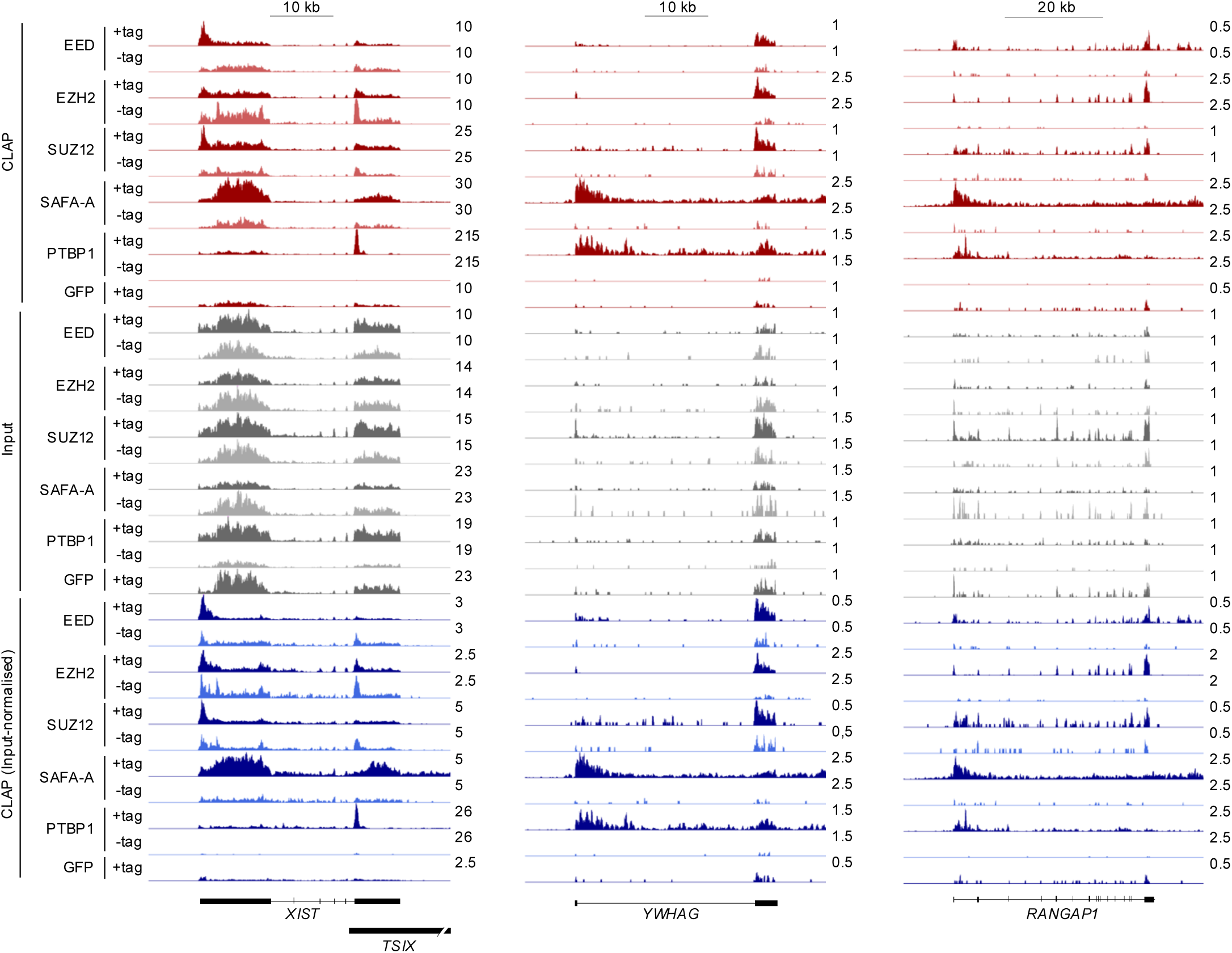
Visualising crosslinked RNA at individual genes supports PRC2 RNA binding in cells. RNA read density (counts per million mapped reads) for +tag and –tag CLAP samples and their respective inputs and for input-normalised CLAP samples at *XI ST*, *YWHAG* and *RANGAP1*. Y-axis scales are equalized between +tag and –tag pairs within a data type. Genes are shown running left to right.

We next examined CLAP signals at individual protein-coding genes. We found many genes for which EED, EZH2 and SUZ12 +tag CLAP samples exhibited stronger signals than GFP and −tag controls, both in terms of raw counts per million and input normalised reads (Fig 4). Thus, consistent with the genome-wide view, examination of the CLAP data at individual genes supported detection of PRC2-RNA interactions in cells.

### RNA segments bound by PRC2 are enriched for G-rich motifs

Multiple methodologies, including CLIP in cells and EMSA and fluorescence anisotropy in vitro, have demonstrated that PRC2 preferentially interacts with G-rich RNA sequences that can form G-quadruplex structures. To determine if PRC2-bound RNA regions identified by CLAP contain these sequences, we used HOMER to identify motifs within RNA windows enriched by CLAP for each of the proteins (Fig 5).

**Fig. 5.**
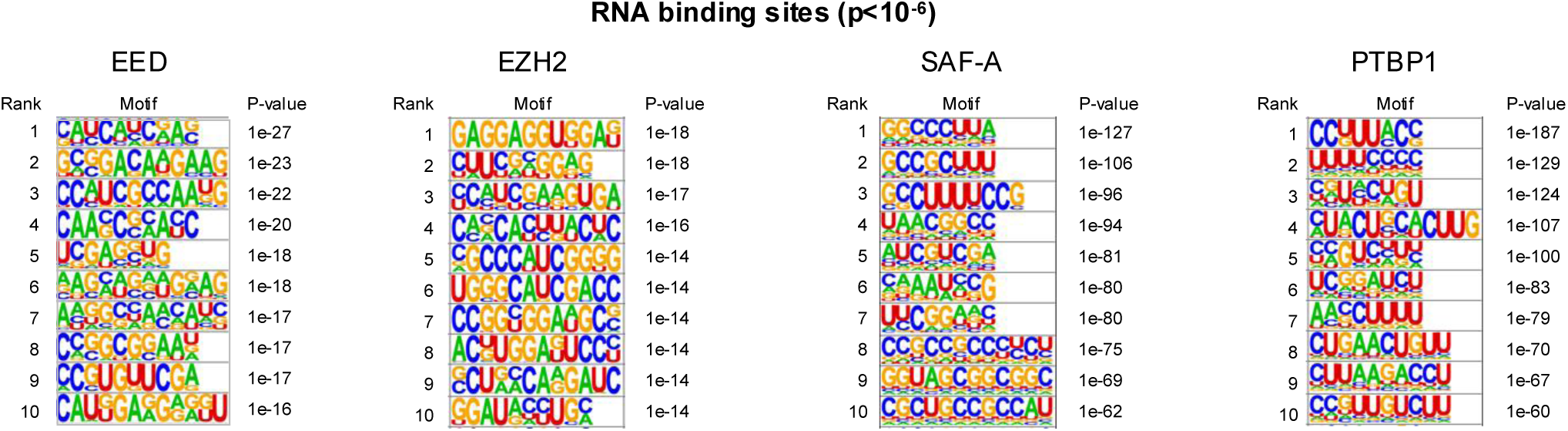
RNA segments bound by PRC2 are enriched for G-rich motifs. The 10 most enriched motifs within the human RNA windows bound by each protein at p<10^-6^ and the significance of their enrichment.

We first examined the motifs enriched (p<10^-6^) in RNA regions bound by SAF-A and PTBP1. RGG/RG amino acids within SAF-A exhibit a preference for binding G4 structures but this selectivity is moderated in the context of the complete protein, which lacks sequence specificity ^20^. Accordingly, we observed various types of motifs enriched at SAF-A-bound RNA regions. In contrast, PTBP1 preferentially binds pyrimidine tracts ^21,22^ and, consistent with this, motifs enriched in PTBP1-bound RNA regions were CU-rich. Next, examining motifs enriched in EZH2-bound RNA regions, we found that the top motif, in addition to a number of others, comprised multiple GG dinucleotides, which allow G4 formation. There was not sufficient power to identify motifs enriched within the small number of SUZ12-bound RNA windows. We conclude that the RNA motifs enriched at EZH2 RNA binding sites identified by CLAP are consistent with the known RNA binding preference of PRC2 ^2,4,6,13^.

## DISCUSSION

Our analysis of the CLAP data generated by Guo and colleagues is consistent with PRC2 RNA binding in living cells. CLAP methodology covalently crosslinks proteins to interacting RNAs and then uses stringent denaturing conditions to purify the tagged proteins of interest and the RNAs to which they are bound. Mixing cells in which the protein is tagged with cells of a second species in which the protein is unmodified provides the ability to differentiate RNA binding in cells from experimental artifacts: CLAP for proteins that bind RNA in cells should show greater enrichment of RNA from the tagged cells versus the unmodified cells of the other species. By this definition, PRC2 is an RNA binding protein. Human RNAs were enriched in EZH2 and, to a lesser extent, EED +tag CLAP samples compared to input, GFP control and - tag samples. The opposite was the case for mouse RNAs, which were enriched in EED and EZH2 −tag CLAP samples compared to +tag samples (albeit CLAP in mouse cells was less efficient). The enrichment of RNAs corresponding to the species in which the proteins were tagged was apparent following the statistical framework presented by Guo et al and using standard bioinformatics tools. We observed greater enrichment of RNA in EZH2 than EED and SUZ12 CLAP experiments, potentially consistent with structural data showing contacts between this subunit and RNA ^13^. Although not as marked as some other CLIP datasets, RNA regions bound by EZH2 were enriched for G-rich motifs, consistent with the RNA binding preference of PRC2 ^2,3,4,6^.

The reason Guo and colleagues did not observe enrichment of RNA in their PRC2 CLAP data appears to be due to their scaling the data by the proportion of genome-mapped reads. This step is not included in the authors’ rationalisation of their data normalisation strategy ^18^ and thus the justification is unclear. Low-input RNA samples produce libraries with low complexity (an over-abundance of rRNA) because stochastic sampling disproportionately captures abundant RNAs and the additional PCR cycles that are often employed in library generation can exacerbate this effect. Ideally, these technical biases are controlled experimentally by spiking RNAs at known frequencies and including unique molecular identifiers (UMIs) within adapters. For datasets with low complexity, such as PRC2 CLAP samples, adjusting for these variables would act to reduce the proportion of multi-copy RNA-mapped reads, thereby increasing the enrichment scores for single-copy RNA windows.

Faced with a lack of UMIs, Guo et al removed reads with identical sequences in the genome-mapped portion only (because reads mapping to multi-copy RNAs will always share sequence, even in the absence of PCR duplication). They then divided read counts by the total read number and multiplied by the proportion of genome-mapped reads (thus, dividing read counts by the total read number a second time). This step had the effect of reducing read counts for samples with low input RNA (such as PRC2 CLAP samples) relative to samples with high input RNA (such as input samples and SAF-A and PTBP1 CLAP samples) and thus reduced enrichment ratios for PRC2 subunits.

Lee and colleagues have recently presented their own re-analysis of Guo and colleagues’ CLAP data in which they normalised read counts by the number of genome-mapped reads ^23^. As set out by Guo et al ^18^, not including multi-copy RNAs in this way has the effect of increasing enrichment scores for low complexity samples such as PRC2. Guo and colleagues scaling procedure skews data the other way by decreasing enrichment scores for low complexity samples. We contend that both of these adjustments inappropriately distort the data and its interpretation.

Regardless, questions of which data normalisation method is most appropriate can largely be negated by comparison between CLAP datasets. Being processed in exactly the same way as +tag samples, control CLAP experiments can be argued to represent better models for background RNA frequencies than input samples. Using both Guo’s statistical framework and standard sequence analysis tools, we found that PRC2 +tag CLAP experiments enriched for human RNAs relative to control GFP and −tag CLAP experiments. Human RNAs can also be seen to have higher enrichment scores in PRC2 +tag CLAP samples versus GFP and −tag samples in Guo and colleagues’ original figures (compare Figs S5 and S8 in ^17^).

A second aspect of Guo and colleagues’ data presentation was potentially misleading. When presenting CLAP data for individual RNAs such as XIST, the authors excluded windows for which the significance of enrichment was below a p<10^-6^ threshold. We agree with Lee and colleagues ^23^ that this filtering gives the impression that PRC2 CLAP datasets don’t contain any reads that map to XIST. Our analysis found that the 5’-end of XIST RNA was significantly enriched in EED and SUZ12 +tag CLAP samples versus input. However, lack of enrichment of this region by EZH2 CLAP and the small peaks within this region in control −tag samples prevent firm conclusions regarding PRC2 binding to XIST from this dataset alone.

Although our analysis identified PRC2 RNA binding in cells, detection was not as robust as for the canonical RNA binding proteins SAF-A and PTBP1. This may be because UV-induced RNA crosslinking is less efficient for PRC2 than SAF-A and PTBP1. Weaker RNA crosslinking to PRC2 subunits compared to PTBP1 can be observed in vitro ^7,17^ and may be due to differences in efficiency with which different amino acids and nucleotides crosslink together ^24–26^. Furthermore, compared to SAF-A and PTBP1, a lower proportion of PRC2 molecules may interact with RNA at any a given time in the cell.

In conclusion, we show here that Guo and colleagues’ CLAP methodology detects interaction of PRC2 with RNA in cells: RNA segments were enriched versus input, and versus GFP and –tag controls. Enrichments versus GFP and −tag controls were robust to the exact analysis method used but enrichment versus input was masked by Guo’s approach of scaling the data by the proportion of genome-mapped reads. Our results are consistent with previous experiments demonstrating PRC2 RNA binding in vitro and in cells and will spur ongoing work seeking to determine the biological impact of PRC2 RNA binding.

## ACKNOWLEDGEMENTS

Thanks to Julian König and Kathi Zarnack for providing feedback on our analysis. Work at the Bioinformatics Translational Technology Platform was supported by the CRUK City of London Centre Award [CTRQQR-2021\100004]. A.H.H. was supported by a PhD studentship funded by the CRUK UCL Centre [C416/A18088].

## AUTHOR CONTRIBUTIONS

Conceptualization, R.G.J.; methodology, S.H., L.C., R.G.J.; formal analysis, S.H., L.C. and A.H.H., R.G.J; writing, R.G.J.; supervision, S.M. and R.G.J..; project administration, R.G.J.

## METHODS

### Data alignment and read counting

Scripts used in our analysis are available at https://github.com/UCL-BLIC/PRC2_CLAP. CLAP sequence data were downloaded from the Sequence Read Archive (SRA: SRP484304; Table 1) and processed using an SGE-compatible version of the Guttman lab CLAPAnalysis pipeline (https://github.com/lconde-ucl/CLAPAnalysis_sge) ^17^, slightly modified to use *bedtools intersect* in place of the *FilterBlacklist.jar* script from the original pipeline (https://github.com/GuttmanLab/CLAPAnalysis). Briefly, adapters were removed with *Trim Galore* (https://github.com/FelixKrueger/TrimGalore), and trimmed reads were aligned to combined sequences of human and mouse multi-copy RNAs downloaded from https://nf-co.re/clipseq/1.0.0/) using *Bowtie 2* ^27^. Reads that did not align were subsequently mapped to a combined mouse (mm10) and human (hg38) reference genome using STAR ^28^. Duplicate reads were removed from the STAR-aligned BAM files with *Picard* (https://github.com/broadinstitute/picard) and reads overlapping multi-copy RNAs (https://github.com/GuttmanLab/CLAPAnalysis/blob/main/clap_pipeline/assets/repeats.RNA. combined.hg38.mm10.bed) were removed with *bedtools intersect* ^29^. The filtered, deduplicated STAR BAM files were then merged with their respective *Bowtie 2* BAM files using *samtools*. Combined BAM files from replicate experiments were merged for each protein.

**Table 1.**
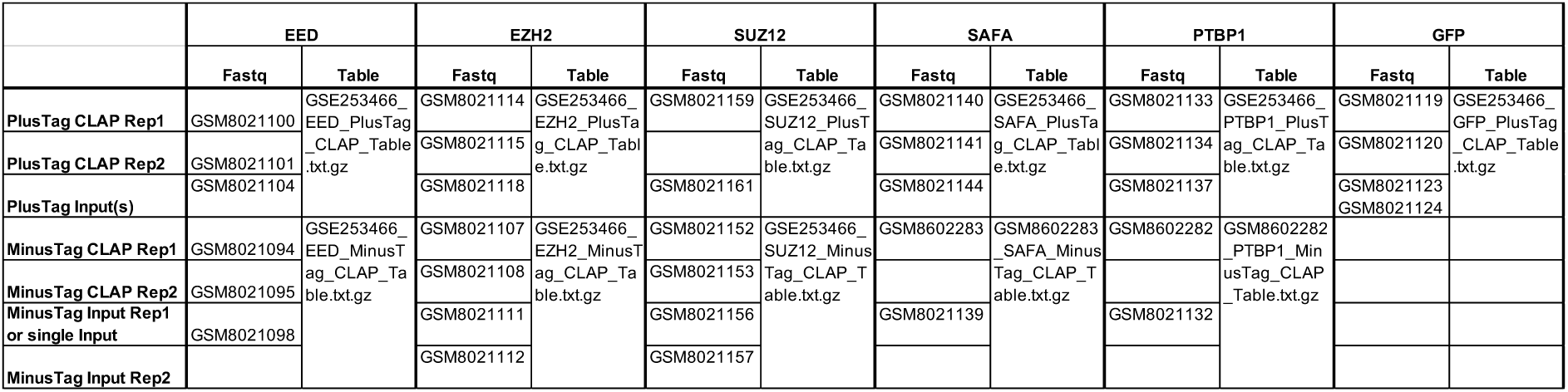
Datasets used in this study. NB. ‘Rep2’ referred to in Fig S8 in Guo et al ^17^ and the supplementary files at GSE253466 do not correspond to the Rep2 fastq files but instead to the “No mixing” samples (personal communication with the authors).

### Enrichment analysis

We first divided UCSC knownCanonical transcripts into 100 bp windows using the start and end gene coordinates from GENCODE gene models (V40 for human and M23 for mouse). Windows were classified as either exon or intron depending on the element with which they exhibited the largest overlap. We recorded the total number of reads (including repetitive and structural RNAs), split the combined BAM files into either human or mouse reads, and then counted reads overlapping each window (1 or more bp) using *bedtools coverage*.

We used an R script (*binomTables.R*) to compare the reads per window of CLAP samples ***C_W_*** to the reads per window of the paired input sample ***I_W_***. The hypothesised probability of success # (expected ratio between the CLAP and input reads) was then calculated as the ratio of all mapped CLAP reads (from the combined BAM files) divided by all mapped reads in the paired input:

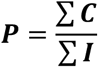

To create a robust floor, the value for each Input window was changed to the maximum of either the number of reads in the window or the median number of reads in all exonic or intronic windows within the same gene (giving ***I_W_*′**). Windows where ***I_W_*′=0** were removed. The enrichment score ***E*** per window ***W*** was then calculated as:

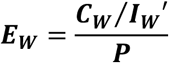

Significance of the enrichment for each window was calculated using the values above (***C_W_*, *I_W_*′, *P*)** with the R *binom.test* function. Data for the transgenes present in each cell line were removed before plotting with ggplot2 in line with Guo et al.

To determine the contribution of variability between input samples to apparent differences between CLAP data sets, we also performed enrichment analysis with single merged +tag and single merged −tag datasets.

### Investigation of data processing by Guo et al

Supplementary data tables containing read counts, enrichment scores and p-values per window calculated by Guo et al were downloaded from GEO (GSE253466; Table 1). The identity of the samples represented by each table was confirmed by Guo et al (personal communication). Normalisation factors used by Guo et al were derived by dividing the window enrichment score by the ratio of the window CLAP read count vs input read count in these tables. Guo and colleagues did not respond to requests to explain the rationale for their data scaling step.

### Differences in data processing to Guo et al

As the multi-copy RNA index used in the Bowtie alignment step was not provided by Guo et al, we instead used the index from the standard nf-core clipseq pipeline (https://nf-co.re/clipseq/1.0.0/). We also tried aligning the data to a multi-copy RNA index generated from the repeat RNA bed file provided by the authors (https://github.com/GuttmanLab/CLAPAnalysis/blob/main/clap_pipeline/assets/repeats.RNA.combined.hg38.mm10.bed), but this resulted in fewer reads being mapped to multi-copy RNAs and did not make a difference to the enrichment of RNAs in PRC2 CLAP experiments. As Guo et al did not provide the coordinates for the 100 nt RNA windows that they defined, we cannot be certain that the RNA windows used here are identical but read count profiles derived from the two sets of windows were highly similar (Fig 2C).

### Plotting enrichment across mature spliced XIST RNA

Exonic windows within XIST were plotted in order from 5’ to 3’-end of the gene and enrichment scores (***E_W_***) plotted.

### Genome browser visualisation

BedGraphs for CLAP and input samples were generated from the combined BAM files using *bamCoverage* with the following options: --binSize 100, --normaliseUsing CPM, --effectiveGenomeSize 5273117239, --extendReads. CLAP sample bedgraphs normalised to their matched input were generated using *bamCompare* with the following options: --binSize 100, --normaliseUsing CPM --scaleFactorsMethod None, --effectiveGenomeSize, 5273117239, --extendReads, --operation ratio, --pseudocount 0 1. The addition of pseudocounts to the input samples served to prevent artificially high CLAP enrichment values caused by normalisation to very low stochastic input values. BedGraph files were then filtered to remove reads mapping to repetitive and structural RNAs (aligned by *Bowtie 2*), split into human and mouse, converted to bigwig files with *bedGraphToBigWig*, and uploaded to the UCSC genome browser. Y-axis scales were equalized between +tag and –tag pairs within a data type.

### Quantifying RNA binding to individual exons and introns

Exonic and intronic genomic coordinates from UCSC knownCanonical transcripts were extracted based on GENCODE gene model files. For each sample, the input-normalised bigWig files generated above were processed with *bigWigAverageOverBed* to calculate the average signal (CPM per base) across each exonic and intronic segment. As for Guo and colleague’s method, exons and introns corresponding to the transgenes were removed. The resulting values were plotted as scatterplots comparing the +tag and –tag conditions for each protein.

### Motif analysis

Motifs enriched within significantly enriched human RNA windows were identified with HOMER with the settings -rna -size given. A random set of 100,000 windows that were not significantly enriched in any of the +tag CLAP experiments was used as the background. The top 10 most significantly enriched motifs were presented.

---

**Fig. S1.**
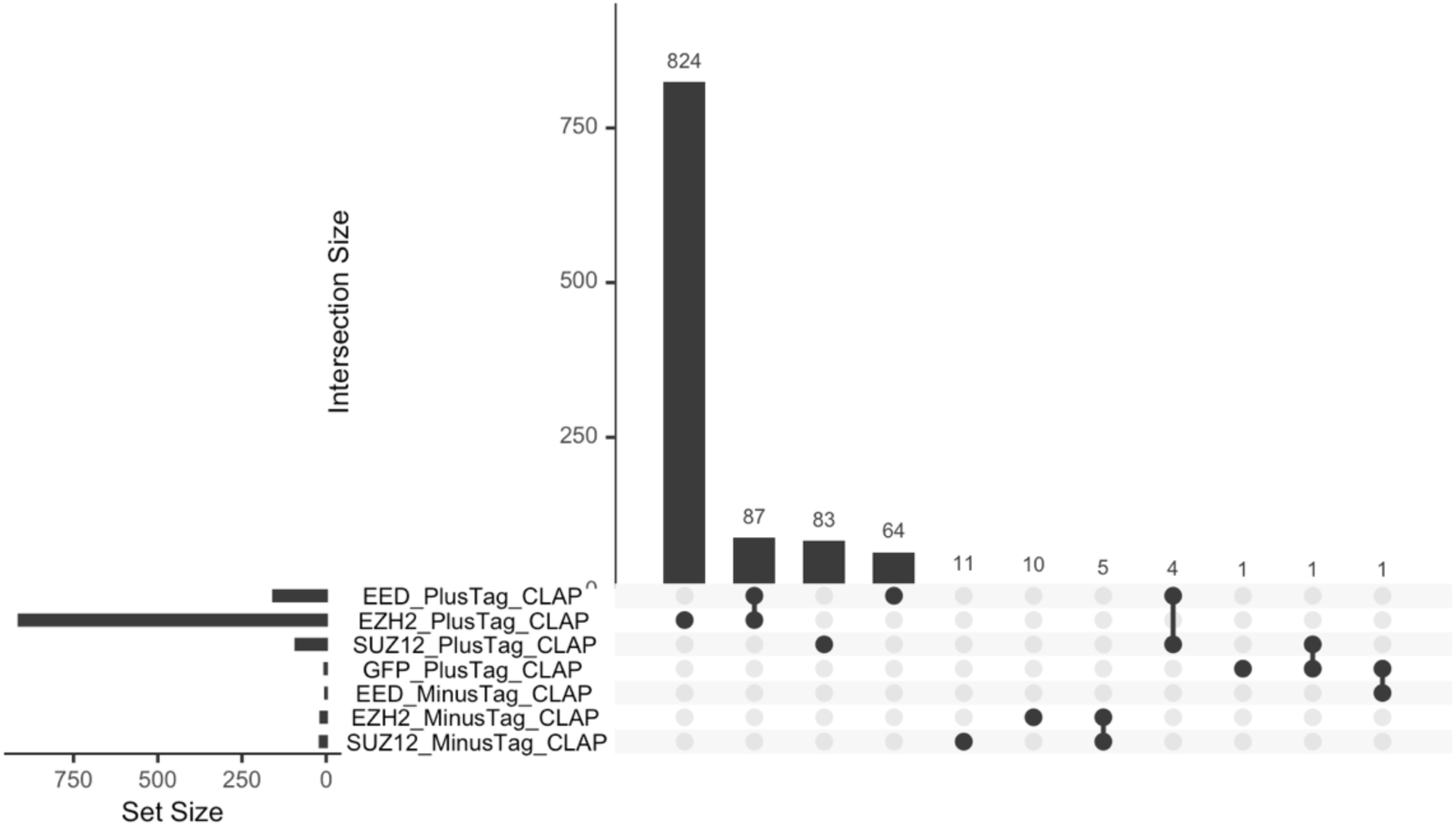
Overlap between significantly enriched RNA windows. UpSet plot showing the intersections between the significantly enriched (p<10 ^-6^) human RNA windows for EED, EZH2, SUZ12 and GFP +tag and –tag CLAP samples.

**Fig. S2.**
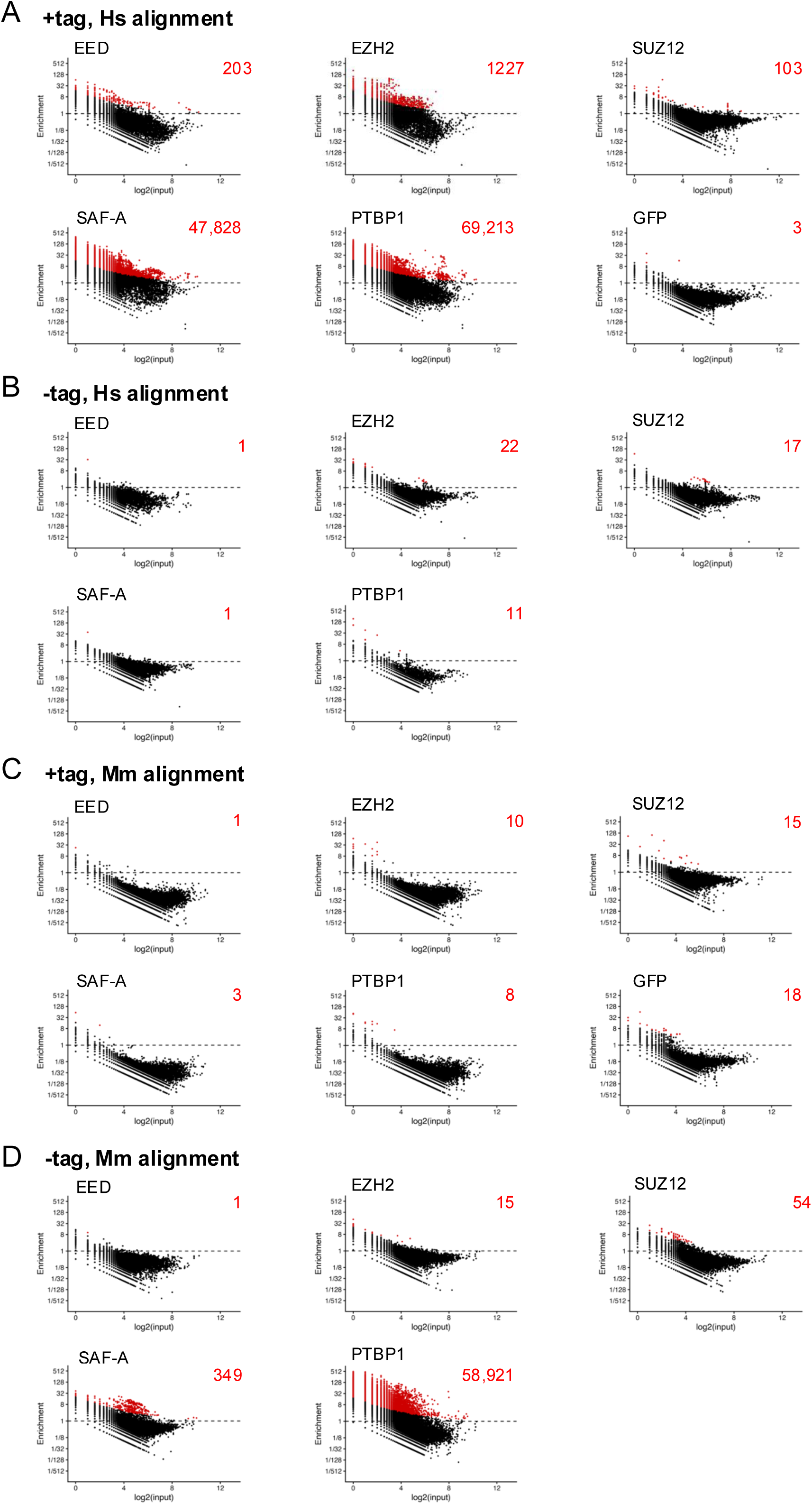
Enrichment of RNA windows at a lower threshold of p < 10^-5^. A. Scatterplots of CLAP enrichment ratio versus paired input read count (both log values) for +tag samples within 100nt windows across human genes. Windows with significant enrichment (p <10^−5^; binomial test) are shown in red and enumerated in the top right of each plot. B. As A., except for −tag CLAP control samples. C. As A., except for +tag CLAP samples and mouse genes. D. As A., except for −tag CLAP samples and mouse genes.

**Fig. S3.**
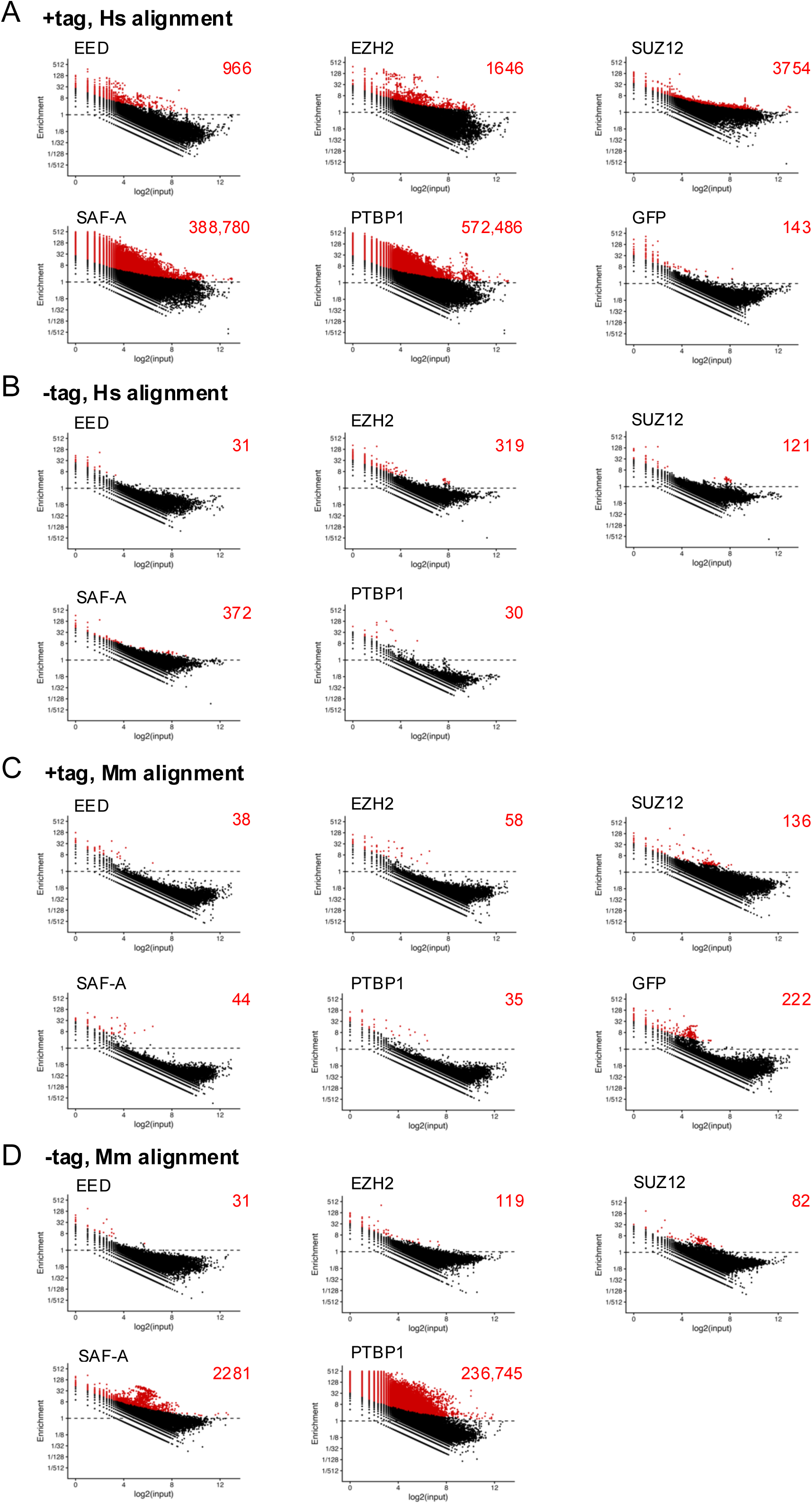
Enrichment of RNA windows relative to a single combined +tag input sample or a combined –tag input sample. A. Scatterplots of +tag CLAP enrichment ratio versus a single combined +tag input read count (both log values) within 100nt windows across human genes. Windows with significant enrichment (p <10^−6^; binomial test) are shown in red and enumerated in the top right of each plot. B. As A., except for −tag CLAP control samples. C. As A., except for +tag CLAP samples and mouse genes. D. As A., except for −tag CLAP samples and mouse genes.

**Fig. S4.**
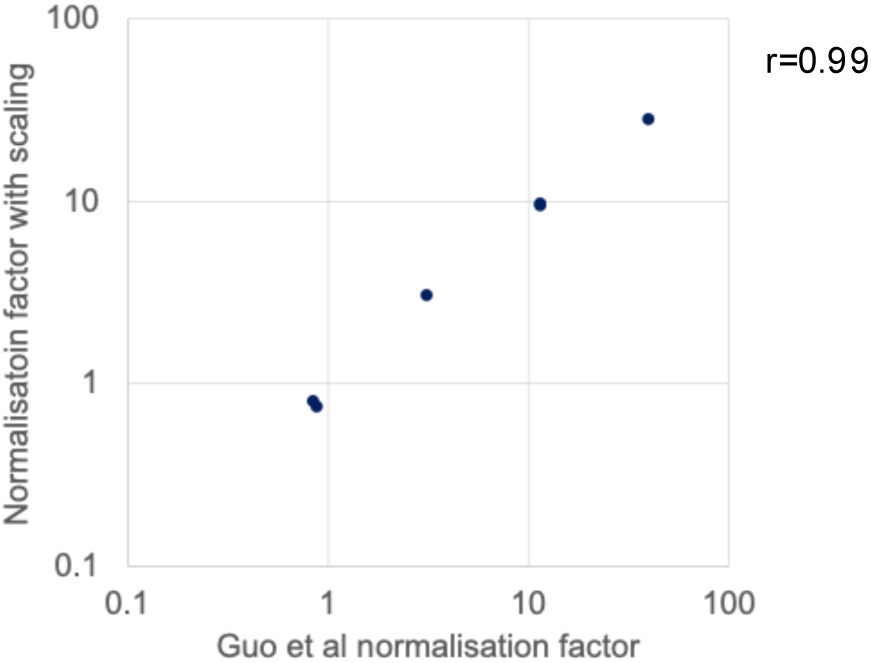
Comparison between normalisation factors used by Guo et al and normalisation factors calculated here by scaling by the proportion of genome-mapped reads. Scatterplot of normalisation factors applied by Guo et al (x-axis, column 4 in Fig 2E) with the normalisation factors calculated here by scaling by the proportion of genome-mapped reads (y-axis, column 5 in Fig 2E). Pearson correlation coefficient = 0.99.

**Fig. S5.**
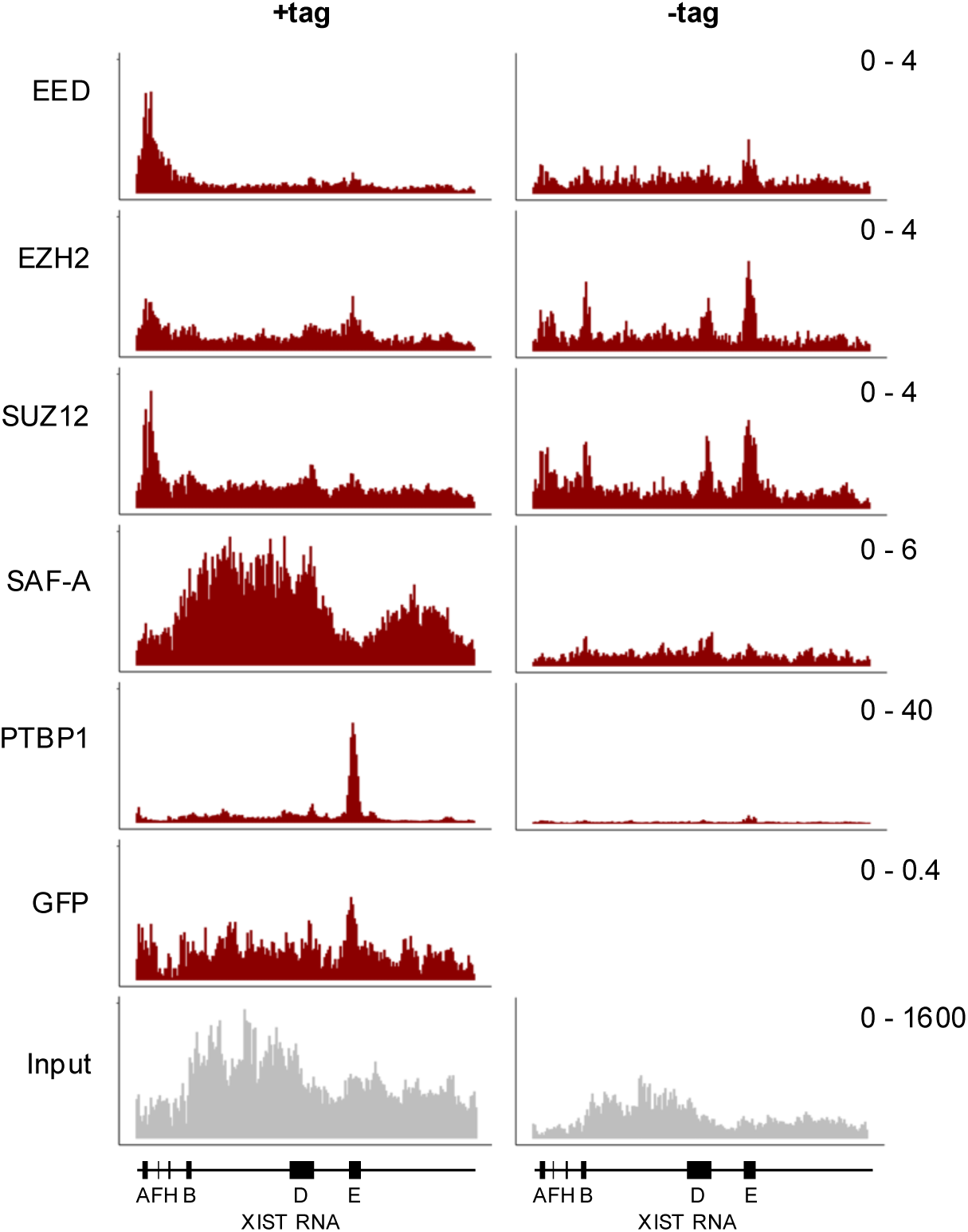
Visualizing PRC2 crosslinking to XIST RNA. Fold enrichment of EED, EZH2, SUZ12, SAF-A, PTBP1 and GFP +tag and –tag CLAP versus their respective inputs, and raw merged +tag input read counts, across human XIST RNA, visualized according to the methodology of Guo and colleagues. Y-axis scales are equalized between +tag and –tag pairs. Repeat regions within XIST are marked. The peaks of EED and SUZ12 binding at the 5’ end of XIST RNA are significant at p<10^-6^.

## REFERENCES

1. Tamburri S, Rustichelli S, Amato S, & Pasini D. Navigating the complexity of Polycomb repression: Enzymatic cores and regulatory modules. Mol Cell. 2024; 84:3381–3405

2. Kaneko S, Son J, Bonasio R, Shen SS, & Reinberg D. Nascent RNA interaction keeps PRC2 activity poised and in check. Genes Dev. 2014; 28:1983–8

3. Wang X, Goodrich KJ, Gooding AR, Naeem H, Archer S, Paucek RD, Youmans DT, Cech TR, & Davidovich C. Targeting of Polycomb Repressive Complex 2 to RNA by Short Repeats of Consecutive Guanines. Mol Cell. 2017; 65:1056–1067

4. Beltran M, Tavares M, Justin N, Khandelwal G, Ambrose J, Foster BM, Worlock KB, Tvardovskiy A, Kunzelmann S, Herrero J, et al. G-tract RNA removes Polycomb repressive complex 2 from genes. Nat Struct Mol Biol. 2019; 26:899–909

5. Zhang Q, McKenzie NJ, Warneford-Thomson R, Gail EH, Flanigan SF, Owen BM, Lauman R, Levina V, Garcia BA, Schittenhelm RB, et al. RNA exploits an exposed regulatory site to inhibit the enzymatic activity of PRC2. Nat Struct Mol Biol. 2019; 26:237–247

6. Rosenberg M, Blum R, Kesner B, Aeby E, Garant JM, Szanto A, & Lee JT. Motif-driven interactions between RNA and PRC2 are rheostats that regulate transcription elongation. Nat Struct Mol Biol. 2021; 28:103–117

7. Song J, Yao L, Gooding AR, Thron V, Hemphill WO, Goodrich K, Erbse AH, Kasinath V, & Cech TR. RNA-induced PRC2 inhibition depends on the sequence of bound RNA. Nat Commun (2026). 10.1038/s41467-026-72294-y

8. Kaneko S, Son J, Shen SS, Reinberg D, & Bonasio R. PRC2 binds active promoters and contacts nascent RNAs in embryonic stem cells. Nat Struct Mol Biol. 2013; 20:1258–64

9. Beltran M, Yates CM, Skalska L, Dawson M, Reis FP, Viiri K, Fisher CL, Sibley CR, Foster BM, Bartke T, et al. The interaction of PRC2 with RNA or chromatin is mutually antagonistic. Genome Res. 2016; 26:896–907

10. Cifuentes-Rojas C, Hernandez AJ, Sarma K, & Lee JT. Regulatory interactions between RNA and polycomb repressive complex 2. Mol Cell. 2014; 55:171–85

11. Herzog VA, Lempradl A, Trupke J, Okulski H, Altmutter C, Ruge F, Boidol B, Kubicek S, Schmauss G, Aumayr K, et al. A strand-specific switch in noncoding transcription switches the function of a Polycomb/Trithorax response element. Nat Genet. 2014; 46:973–981

12. Wang X, Paucek RD, Gooding AR, Brown ZZ, Ge EJ, Muir TW, & Cech TR. Molecular analysis of PRC2 recruitment to DNA in chromatin and its inhibition by RNA. Nat Struct Mol Biol. 2017; 4:1028–1038

13. Song J, Gooding AR, Hemphill WO, Love BD, Robertson A, Yao L, Zon LI, North TE, Kasinath V, & Cech TR. Structural basis for inactivation of PRC2 by G-quadruplex RNA. Science. 2023; 381:1331–1337

14. Skalska L, Begley V, Beltran M, Lukauskas S, Khandelwal G, Faull P, Bhamra A, Tavares M, Wellman R, Tvardovskiy A, et al. Nascent RNA antagonizes the interaction of a set of regulatory proteins with chromatin. Mol Cell. 2021; 81:2944–2959

15. Long Y, Hwang T, Gooding AR, Goodrich KJ, Rinn JL, & Cech TR. RNA is essential for PRC2 chromatin occupancy and function in human pluripotent stem cells. Nat Genet. 2020; 52:931–938

16. Gail EH, Healy E, Flanigan SF, Jones N, Ng XH, Uckelmann M, Levina V, Zhang Q, & Davidovich C. Inseparable RNA binding and chromatin modification activities of a nucleosome-interacting surface in EZH2. Nat Genet. 2024; 56:1193–1202

17. Guo JK, Blanco MR, Walkup WGt, Bonesteele G, Urbinati CR, Banerjee AK, Chow A, Ettlin O, Strehle M, Peyda P, et al. Denaturing purifications demonstrate that PRC2 and other widely reported chromatin proteins do not appear to bind directly to RNA in vivo. Mol Cell. 2024; 84:1271–1289

18. Guo JK, Blanco MR, & Guttman M. Failing to account for RNA quantity inflates background and leads to the misleading appearance that PRC2 and GFP bind to RNA in vivo. bioRxiv 2024; 10.1101/2024.11.02.621417

19. Cech TR, Davidovich C, & Jenner RG. PRC2-RNA interactions: Viewpoint from Tom Cech, Chen Davidovich, and Richard Jenner. Mol Cell. 2024; 84:3593–3595

20. Kletzien OA, Wuttke DS, & Batey RT. The RNA-binding Selectivity of the RGG/RG Motifs of hnRNP U is Abolished by Elements Within the C-terminal Intrinsically Disordered Region. J Mol Biol. 2024; 436:168702

21. Perez I, Lin CH, McAfee JG, & Patton JG. Mutation of PTB binding sites causes misregulation of alternative 3’ splice site selection in vivo. RNA. 1997; 3:764–78

22. Singh R, Valcarcel J, & Green MR. Distinct binding specificities and functions of higher eukaryotic polypyrimidine tract-binding proteins. Science. 1995; 268:1173–6

23. Lee Y, Das P, Kesner B, Rosenberg M, Blum R, & Lee JT. Re-analysis of CLAP data affirms PRC2 as an RNA binding protein. bioRxiv 2024; 10.1101/2024.09.19.613009

24. Knorlein A, Sarnowski CP, de Vries T, Stoltz M, Gotze M, Aebersold R, Allain FH, Leitner A, & Hall J. Nucleotide-amino acid pi-stacking interactions initiate photo cross-linking in RNA-protein complexes. Nat Commun. 2022; 13:2719

25. McGregor A, Rao MV, Duckworth G, Stockley PG, & Connolly BA. Preparation of oligoribonucleotides containing 4-thiouridine using Fpmp chemistry. Photo-crosslinking to RNA binding proteins using 350 nm irradiation. Nucleic Acids Res. 1996; 24:3173–80

26. Feng H, Lu X, Maji S, Liu L, Ustianenko D, Rudnick N, & Chaolin Z. Structure-based prediction and characterization of photo-crosslinking in native protein-RNA complexes. Nat Commun. 2024; 15:2279.

27. Langmead B and Salzberg SL. Fast gapped-read alignment with Bowtie 2. Nat Methods. 2012; 9:357–9

28. Dobin A, Davis CA, Schlesinger F, Drenkow J, Zaleski C, Jha S, Batut P, Chaisson M, & Gingeras TR. STAR: ultrafast universal RNA-seq aligner. Bioinformatics. 2013l 29:15–21

29. Quinlan AR and Hall IM. BEDTools: a flexible suite of utilities for comparing genomic features. Bioinformatics. 2010; 26:841–2

